# Nonuniform scaling of synaptic inhibition in the dorsolateral geniculate nucleus in a mouse model of glaucoma

**DOI:** 10.1101/2024.03.27.587036

**Authors:** Matthew J. Van Hook, Shaylah McCool

## Abstract

Elevated intraocular pressure (IOP) triggers glaucoma by damaging the output neurons of the retina called retinal ganglion cells (RGCs). This leads to the loss of RGC signaling to visual centers of the brain such as the dorsolateral geniculate nucleus (dLGN), which is critical for processing and relaying information to the cortex for conscious vision. In response to altered levels of activity or synaptic input, neurons can homeostatically modulate postsynaptic neurotransmitter receptor numbers, allowing them to scale their synaptic responses to stabilize spike output. While prior work has indicated unaltered glutamate receptor properties in the glaucomatous dLGN, it is unknown whether glaucoma impacts dLGN inhibition. Here, using DBA/2J mice, which develop elevated IOP beginning at 6-7 months of age, we tested whether the strength of inhibitory synapses on dLGN thalamocortical relay neurons is altered in response to the disease state. We found an enhancement of feed-forward disynaptic inhibition arising from local interneurons along with increased amplitude of quantal inhibitory synaptic currents. A combination of immunofluorescence staining for the GABA_A_-α1 receptor subunit, peak-scaled nonstationary fluctuation analysis, and measures of homeostatic synaptic scaling indicated this was the result of an approximately 1.4-fold increase in GABA receptor number at post-synaptic inhibitory synapses, although several pieces of evidence strongly indicate a non-uniform scaling across inhibitory synapses within individual relay neurons. Together, these results indicate an increase in inhibitory synaptic strength in the glaucomatous dLGN, potentially pointing toward homeostatic compensation for disruptions in network and neuronal function triggered by increased IOP.

**Significance Statement:** Elevated eye pressure in glaucoma leads to loss of retinal outputs to the dorsolateral geniculate nucleus (dLGN), which is critical for relaying information to the cortex for conscious vision. Alterations in neuronal activity, as could arise from excitatory synapse loss, can trigger homeostatic adaptations to synaptic function that attempt to maintain activity within a meaningful dynamic range, although whether this occurs uniformly at all synapses within a given neuron or is a non-uniform process is debated. Here, using a mouse model of glaucoma, we show that dLGN inhibitory synapses undergo non-uniform upregulation due to addition of post-synaptic GABA receptors. This is likely to be a neuronal adaptation to glaucomatous pathology in an important sub-cortical visual center.

## Introduction

The goal of this study was to determine whether glaucoma, a disease that culminates in blindness from loss of retinal ganglion cells (RGCs) and their projections to the brain (Calkins, 2012; Weinreb et al., 2014), leads to homeostatic scaling of inhibitory synapses made onto thalamocortical (TC) relay neurons in the dorsolateral geniculate nucleus (dLGN) of the thalamus. The dLGN processes visual inputs directly from RGCs and transmits that information to the cortex, making it critical for conscious vision (Kerschensteiner and Guido, 2017; Guido, 2018; Liang and Chen, 2020). dLGN TC neurons receive inhibition from two principal sources - feedback projections from the thalamic reticular nucleus (TRN) and feedforward inhibition from local inhibitory interneurons (LINs) - that play important roles in shaping TC neuron receptive fields and setting resting membrane potential and firing mode (Wang et al., 2011; Hirsch et al., 2015; Cox and Beatty, 2017).

Synaptic scaling is a homeostatic neuronal adaptation whereby altered synaptic inputs and/or disrupted activity levels lead neurons to modulate the synaptic strength to maintain action potential output (Turrigiano et al., 1998; Turrigiano, 2012). This phenomenon has been reported to occur at both inhibitory and excitatory synapses in response to activity modulation in dissociated cell cultures, in the visual cortex in vivo through intraocular TTX injection or visual deprivation, in the auditory system following hearing loss, and in response to seizure activity (Otis et al., 1994; Turrigiano et al., 1998; Desai et al., 2002; Wierenga et al., 2005; Ibata et al., 2008; Maffei and Turrigiano, 2008; Goold and Nicoll, 2010; Joseph and Turrigiano, 2017; Teichert et al., 2017; Gonzalez-Islas et al., 2023). Conventionally, synaptic scaling is thought to be the result of a global multiplicative process that triggers uniform changes at each of a neuron’s synaptic inputs via somatic Ca^2+^ signals (Ibata et al., 2008; Goold and Nicoll, 2010; Wenner, 2011; Turrigiano, 2012; Joseph and Turrigiano, 2017). However, it has become increasingly apparent that many neurons employ a “divergent” scaling process whereby scaling is non-uniform, with different synapses within individual neurons undergoing different degrees of scaling (Hanes et al., 2020; Koesters et al., 2024). However, the mechanisms and conditions driving such scaling are still debated.

We have shown previously that during glaucoma, retinal output signaling to the dLGN is disrupted, which is apparent as an intraocular pressure (IOP)-dependent diminishment of RGC excitatory drive to post-synaptic TC neurons (Van Hook et al., 2020; Smith et al., 2022). We have also found that in both a mouse model of glaucoma and in mice where bilateral enucleation leads to a loss of RGC synaptic input to the dLGN, TC neurons display an increase in intrinsic excitability, firing more action potentials in response to depolarizing stimulation (Van Hook et al., 2020; Bhandari et al., 2022; Smith et al., 2022). Although disrupted synaptic function and altered TC neuron spiking seem to be likely potential triggers for homeostatic scaling of excitatory synapses in the dLGN, there were no detectable effects on quantal excitatory current amplitudes in those studies. However, as diminished drive from RGCs is expected to also disrupt other members of the dLGN synaptic network such as feed-forward inhibition, it remains to be seen whether glaucoma alters dLGN inhibitory synapses. Here, we tested this possibility, finding an increase in the strength inhibitory synapses on dLGN TC neurons from 10-14 month-old mice with glaucoma. This was apparent as an approximately 1.4-fold increase in the number of GABA (γ-aminobutyric acid) receptors present at inhibitory synapses without changes in single channel conductance, although several parallel analysis approaches point toward non-uniform “divergent” scaling across the population of inhibitory synapses on individual TC neurons. Together, these results indicate a scaling up of synaptic strength at a population of inhibitory synapses in the dLGN in mice with elevated IOP, possibly pointing to a homeostatic adaptation to disrupted function of the visual pathway accompanying glaucomatous pathology.

## Materials and Methods

### Animals

For this study, we used male and female DBA/2J (D2; Jackson Labs #000671, RRID:IMSR_JAX:000671)(John et al., 1998) and DBA/2J^gpnmb+^ (D2-control; Jackson Labs #007048, RRID:IMSR_JAX:007048) (Howell et al., 2007) mice, aged 10-14 months. All animal procedures were approved by the Institutional Animal Care and Use Committee at the University of Nebraska Medical Center. Mice were housed in a 12/12-hour light/dark cycle and provided food and water *ad libitum*. Intraocular pressure measurements were taken in mice lightly anesthetized with isoflurane (2%) approximately monthly beginning at 2-5 months of age using a TonoLab rebound tonometer. D2 mice (N=43) had significantly higher peak IOPs than the D2-control mice (N = 41), with maximum IOP averages of both eyes 15.6±0.3 mmHg for D2-control mice and 23.9±0.8 mmHg for D2 mice (p=1.3×10^-13^, unpaired t-test).

### Slice preparation and electrophysiology

For patch-clamp recording, we prepared 250 micron-thick coronal slices or 300 micron-thick parasagittal slices (Turner and Salt, 1998; Chen and Regehr, 2000; Smith et al., 2022) containing the dLGN and optic tract using the protected recovery method (Ting et al., 2014, 2018). Mice were killed by CO_2_ asphyxiation and cervical dislocation and quickly decapitated. Brains were dissected into a slush of artificial cerebrospinal fluid (aCSF) comprised of (in mM) 128 NaCl, 2.5 KCl, 1.25 NaH_2_PO_4_, 24 NaHCO_3_, 12.5 glucose, 2 CaCl_2_, and 2 MgSO_4_ and bubbled with 5% CO_2_/95% O_2_. After chilling in aCSF for approximately 30 seconds, brains were hemisected using a razor blade with a cut oriented ∼5 degrees from the medial longitudinal fissure and ∼20 degrees from the horizontal plane. The cut surface was glued to a flat agar block positioned on the stage of a Leica VT1000S vibratome which was subsequently submerged in ice-cold aCSF. 300 micron-thick slices were cut, visually inspected to verify they contained dLGN and optic tract, and submerged in an *N*-methyl-D-glucamine-based solution (in mM: 92 NMDG, 2.5 KCl, 1.25 NaH_2_PO_4_, 25 glucose, 30 NaHCO_3_, 20 HEPES, 0.5 CaCl_2_, 10 MgSO_4_, 2 thiourea, 5 L-ascorbic acid, and 3 Na-pyruvate, bubbled with 5% CO_2_/95% O_2_ and warmed to 33°C) for 12 minutes. After the NMDG solution incubation, slices were transferred to room temperature aCSF and incubated for at least 1 hour prior to initiating patch clamp recording.

For patch clamp recording, slices were transferred to a recording chamber, secured with a nylon net, positioned on the stage of an Olympus BX51WI microscope, and superfused with aCSF warmed to 30-33°C with an in-line solution heater (Van Hook, 2020). For optic tract stimulation in parasagittal slices, a broken patch pipette (opening = ∼50 microns) filled with aCSF was positioned in the optic tract ∼500 microns from the dLGN and used as a stimulating electrode. dLGN thalamocortical (TC) relay neurons were targeted for whole-cell voltage clamp recording based on soma size and morphology using patch pipettes (4-7 MΩ) filled with a Cs-based solution (in mM: 120 Cs-methanesulfonate, 2 EGTA, 10 HEPES, 8 TEA-Cl, 5 ATP-Mg, 0.5 GTP-Na_2_, 5 phosphocreatine-Na_2_, 2 QX-314). Reported voltages are corrected for a 10 mV liquid junction potential. An Axon Multiclamp 700B amplifier and Digidata 1550 digitizer (Molecular Devices) were used for electrophysiology data acquisition. Current stimuli (0.1-0.3 ms, up to 1 mA) were delivered to the optic tract using an A-M Systems Instruments Isolated Pulse Stimulator. Signals were sampled on-line at 10 kHz. For each cell, stimulus amplitude, sign, and duration were adjusted to the minimum stimulus strength needed to reliably obtain a maximal excitatory response amplitude measured at -70 mV. Feed-forward disynaptic inhibition (Blitz and Regehr, 2005; Augustinaite and Heggelund, 2018) was measured in response to optic tract stimulation by voltage clamping at 0 mV. Series resistance was partially compensated (60-70%) using the amplifier circuitry. When used, as indicated in the Results section, pharmacological agents (CNQX, D-AP5, TTX, SR95531) were added to the superfusate from a 1000x stock solutions and bath-applied. CNQX, D-AP5, and SR95531 (Gabazine) were purchased from Tocris while tetrodotoxin was purchased from Abcam. CNQX and SR95531 stocks were prepared in dimethylfulfoxide, while D-AP5 and TTX stocks were prepared in deionized water.

Miniature inhibitory post-synaptic currents (mIPSCs) were measured at 0 mV in the absence of stimulation using aCSF supplemented with 20 μM CNQX, 50 μM D-AP5, and 0.5-1 μM tetrodotoxin (TTX). No series resistance compensation was used for mIPSC recordings due to the increased noise it contributes. mIPSCs were detected and analyzed using MiniAnalysis software (Synaptosoft) with an amplitude threshold of 4.5 pA. Fits of mIPSC amplitude and baseline noise distributions revealed good separation of recording noise and mIPSCs with these detection parameters. mIPSC amplitude for individual cells is reported as the average amplitude of all events analyzed from that cell. Reported mIPSC frequency for each cell is the median instantaneous frequency of analyzed events from that cell.

We used two analysis approaches to test for and determine the degree of inhibitory synaptic scaling from mIPSC data. In the first, we used linear regression to fit the relationship of rank-ordered mIPSC amplitudes from D2-control and D2 TC neurons (Turrigiano et al., 1998; Hanes et al., 2020). For this, 1900 mIPSCs from D2 recordings and 1900 mIPSCs from D2-control recordings were randomly selected to ensure an equal number of mIPSCs from each cell from a genotype along with an equal overall number of events between D2 and D2-control. Based on the slope (scaling factor) of the best-fit line to the rank-order matched scatterplot, we down-scaled the D2 mIPSCs, excluded those falling below the mIPSC detection threshold (4.5 pA), and compared the cumulative distribution of down-scaled mIPSCs with the control mIPSC distribution using a Komolgorov-Smirnov (K-S) test. In the second approach (Kim et al., 2012), we iteratively down-scaled the population of D2 mIPSC amplitudes using scaling factors of 1.05 to 1.85. After pruning down-scaled event amplitudes that fell below our detection threshold (4.5 pA), we compared the cumulative distributions of D2-control and down-scaled D2 mIPSC amplitudes using a Komolgorov-Smirnov test, using finer gradations of scaling factors to determine the one that produced the highest K-S p-value.

### Peak-scaled nonstationary fluctuation analysis

To estimate single channel conductance and numbers of synaptic GABA_A_ receptors in dLGN TC neurons, we performed a peak-scaled nonstationary fluctuation analysis of the decay phase of mIPSCs (Traynelis et al., 1993; Kilman et al., 2002). In cells with low enough mIPSC frequency to ensure no overlap of the mIPSC decay with other events, we scaled the average peak-aligned mIPSC waveform to the peak amplitude of each event (18-68 events per cell) and calculated the variance over the decay phase. Cells were included in this analysis if there was no statistically significant linear correlation of mIPSC peak amplitude with decay time constant, indicating adequate voltage clamp. The relationship of variance to the mean amplitude during the IPSC decay was fit with a parabolic function:

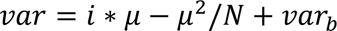

where *var* is the variance, *i* is the single channel current, *µ* is the mean current, *N* is the number of open GABA_A_ receptors at the mIPSC peak, and *var_b_* is the baseline variance. Single channel conductance (γ) was calculated from the fit parameter *i* using the Cl^-^ driving force of 70 mV.

### Histology

For immunofluorescence staining for the vesicular GABA transporter (vGAT) or GABA_A_-α1 GABA receptor subunit in fixed brain sections, mice were killed by CO_2_ asphyxiation and cervical dislocation. Brains were rinsed briefly in phosphate buffered saline (PBS) and then immersed in 4% paraformaldehyde in PBS for 4 hours while rocking at room temperature. Brains were then rinsed 3x 15 minutes with PBS and immersed in 30% sucrose in PBS overnight for cryoprotection. Brains were then embedded in 3% agar and 50 micron-thick sections were cut using a Leica VT1000S vibratome and mounted on Superfrost Plus slides, which were stored at - 20°C. For immunofluorescence staining, slices were thawed and rinsed briefly in PBS. Mounted brain sections were blocked and permeabilized for 1 hour at room temperature in a blocking buffer comprised of 0.5% TritonX-100 and 5.5% donkey serum in PBS. They were then incubated for two nights in primary antibody (1:500 Rabbit-anti-GABA_A_-α1; Synaptic Systems 224 203, RRID:AB_2232180 or Rabbit-anti-vGAT, Synaptic Systems 131 002, RRID:AB_887871) at 4°C, rinsed 6×10 mins in PBS, incubated 2.5 hours in an AlexaFluor-568 conjugated secondary antibody (1:200 donkey-anti-rabbit IgG conjugated to AlexaFluor 568, ThermoFisher A10042, RRID:AB_2534017), rinsed 3×5 mins, and coverslipped with VectaShield Hardset.

Imaging was performed using a Scientifica SliceScope equipped with 2-photon imaging capabilities including a MaiTai HP Ti:sapphire laser that was tuned to 800 nm with power set to 80 μW. A series of optical sections were acquired through the z-axis of the tissue (0.3 μm spacing) with a 74×74 μm field of view (14 pixels/micron). Four images were acquired at each optical plane and averaged to reduce noise. For analysis, background was subtracted using a 50 pixel rolling ball for GABA_A_-α1. For GABA_A_-α1, the average pixel intensity of each image in the stack was measured and the average intensity values of the six peak optical sections through the image stack was used for the intensity. vGAT images were filtered with the “unsharp mask” tool (25 pixel radius, weight = 0.6). vGAT puncta were detected and analyzed and regions of interest generated for each punctum using the Synapse Counter plug-in with a 10 μm rolling ball radius and a min-max size threshold of 0.26-39 μm^2^.

### Experimental Design and Statistical Analysis

Statistical analysis was performed using GraphPad 10. Normality of data was assessed using a D’Agostino & Pearson test. Significant differences between normally distributed datasets were assessed using a two-tailed nested t-test for instances where we recorded multiple cells for individual mice. This approach avoids pitfalls arising from pseudoreplication (Eisner, 2021). Datasets that followed a logarithmic distribution were log transformed prior to statistical testing. For instances where we made single measurements from individual mice (immunofluorescence staining), we used a conventional unpaired t-test. A paired t-test was used to test for effects of acute TTX application on individual cells. Data from individual cells are displayed as individual points for electrophysiology analyses and for individual mice for immunofluorescence analysis. A Komolgorov-Smirnov (K-S) test was used to compare cumulative distributions. An alpha value of 0.05 was selected as the threshold for statistical significance for all tests. For vGAT analysis, the intensity of each vGAT punctum was measured as the average pixel intensity within regions of interest corresponding to each detected punctum. vGAT density was calculated for each image and D2 and D2-control measurements compared using a t-test. The vGAT punctum size and intensity were log transformed before comparison with a nested t-test. Data are presented as mean±SEM for normally distributed data and median(inter-quartile range) for non-normally distributed data, as indicated in the text. Sample sizes (number of mice and number of cells) are reported in the figure legends.

## Results

To test the possibility that glaucoma leads to homeostatic alterations at feed-forward inhibitory synapses in the dLGN, we prepared parasagittal slices containing the dLGN and stimulated retinal ganglion cell axons by positioning a stimulating electrode in the optic tract. The current stimulus amplitude was increased to reliably evoke a maximal response. By voltage clamping TC neurons at the chloride and cationic equilibrium potentials (-70 and 0 mV, respectively), we were able to isolate both monosynaptic retinogeniculate excitatory input and disynaptic feed-forward inhibition (Blitz and Regehr, 2005; Augustinaite and Heggelund, 2018), which arises from local inhibitory interneurons (Figure 1A-D). Confirming adequate isolation of inhibitory and excitatory responses, the synaptic current evoked while TC neurons were voltage clamped at 0 mV was entirely blocked by application of SR95531 (20 μM; 100±2% block; n = 6 cells, p=0.027 paired t-test; Figure 1A). Similar to our previous work, the amplitude of the retinogeniculate EPSC was lower in slices from D2 mice compared to D2-controls (median (IQR), D2-control: 1734(1188-2120) pA, n=28 cells, 9 mice; D2: 668(334-1417) pA, n = 39 cells, 9 mice; p=0.0014, nested t-test of log-transformed data), indicating a loss of retinogeniculate input in the D2 mouse dLGN (Figure 1E). The amplitude of the disynaptic IPSC was also somewhat lower in D2 relative to D2 control (D2-control: 856(624-1216) pA; D2: 578(350-851) pA; p=0.039, nested t-test of log-transformed data), pointing to a reduction in the strength of feed-forward inhibition from local interneurons (Figure 1F).

**Figure 1.**
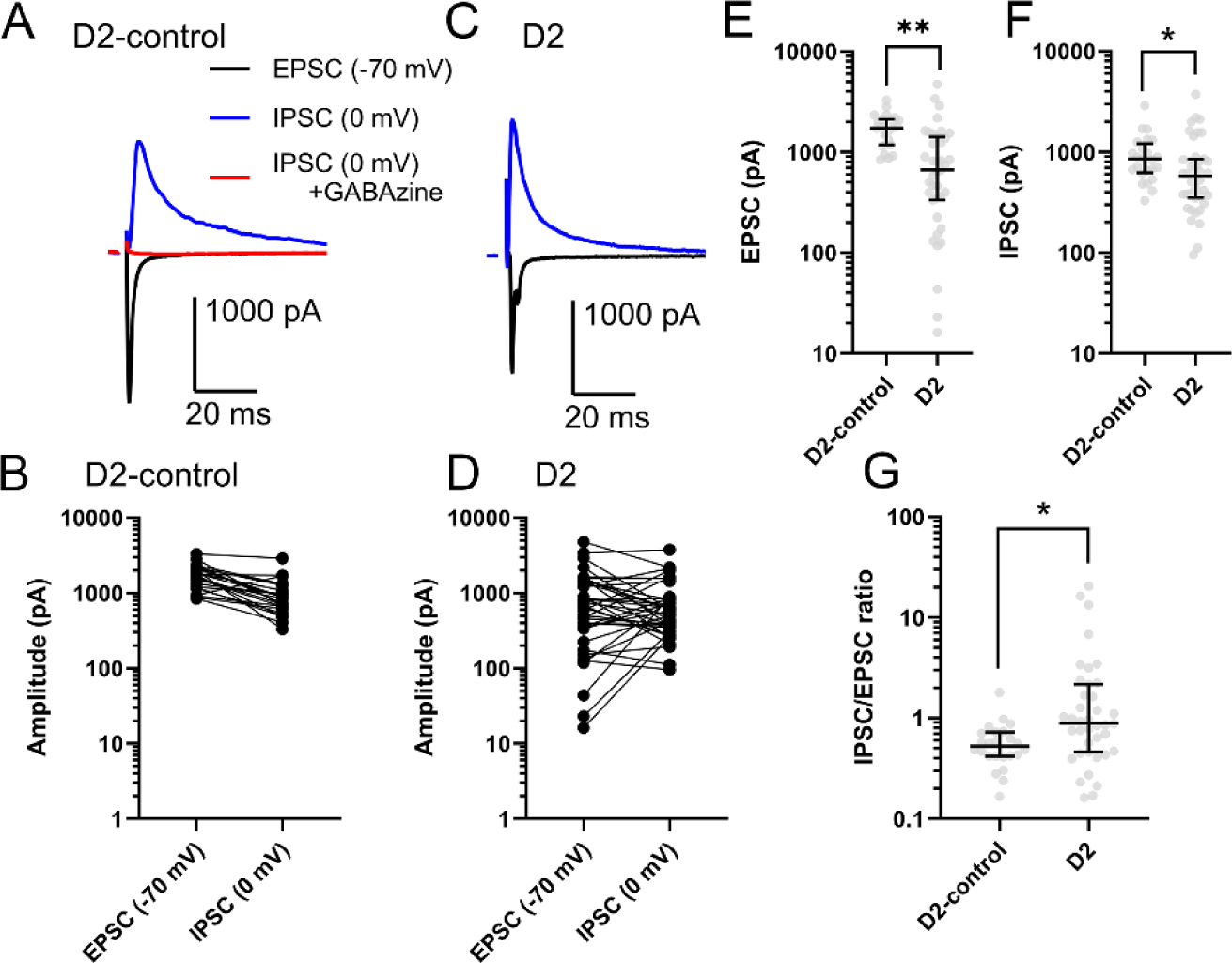
Retinogeniculate excitation and feed-forward disynaptic inhibition in dLGN brain slice recordings from D2-control and D2. **A)** Voltage-clamp recording of monosynaptic retinogeniculate EPSC (black, -70 mV) in response to optic tract stimulation in a brain slice from a D2-control mouse. The disynaptic IPSC was recorded at 0 mV (blue) and was entirely blocked by GABAzine (SR95531, 20 µM, red). **B)** Group data of EPSC and IPSC amplitudes (28 cells from 9 mice). **C)** EPSC and IPSC recordings from a D2 mouse. **D)** Group data of EPSC and IPSC amplitudes from D2 brain slices (39 cells from 9 mice). **E & F)** Group data of EPSC and IPSC amplitudes. Nested t-test performed on log-transformed data. **p<0.01; *p<0.05. **G)** Normalizing the IPSC to the EPSC amplitude shows a relative increase in the strength of feed-forward inhibition in recordings from D2 brain slices. Nested t-test performed on log-transformed data. *p<0.05.

We next sought to determine whether this reduction in IPSC amplitude scaled with the reduced EPSC. This was undertaken to test whether the reduced feed-forward inhibition simply reflects reduced strength of excitatory drive from RGC axons, which would be the case if inhibition was diminished in proportion to the diminishment of excitation. We assessed this by normalizing the IPSC to the EPSC amplitudes for individual cells (IPSC/EPSC; Figure 1G). If that ratio is dissimilar in D2-control vs. D2 recordings, then it would imply that mechanisms downstream of retinogeniculate excitation are acting on the feed-forward inhibitory synapses to modulate their strength. In this analysis the IPSC/EPSC was 0.53(0.42-0.73) [median(IQR)] for D2-control and 0.88(0.46-2.2) for D2 slices (p=0.016, nested t-test of log-transformed data). The higher IPSC/EPSC ratio in TC neurons from D2 mice indicates that feed-forward inhibition, while reduced relative to controls, is higher than would be expected if it simply scaled with the reduced retinogeniculate drive. This implies a relative scaling up of feed-forward inhibition to TC neurons in the D2 dLGN.

To test whether this enhancement of feed-forward inhibition reflects changes in the quantal properties of inhibitory synaptic transmission in the dLGN, we next sought to record single-vesicle synaptic currents in the absence of stimulation. This was accomplished by voltage-clamping TC neurons at 0 mV in the presence of inhibitors of glutamatergic synapses (20 μM CNQX and 50 μM D-AP5). We found that acute application of tetrodotoxin (0.5-1 μM) led to a reduction in both IPSC amplitude (16±2 pA to 12±1 pA, n = 13 cells; p=0.009, paired t-test) and frequency (48±2 Hz to 33±3 Hz, n = 13 cells, p=8×10^-6^, paired t-test), suggesting that even under glutamatergic blockade, spontaneous inhibition represents some level of action potential-driven multiquantal GABA release in the dLGN (Figure 2). Therefore, we performed recordings in the presence of TTX along with glutamatergic blockers to isolate quantal miniature IPSCs (mIPSCs). Under these conditions, we found that the mIPSC frequency was not significantly different between recordings from D2-control and D2 mice (D2-control: 34±2 Hz, n = 19 cells, 8 mice; D2: 31±3 Hz, n = 22 cells, 9 mice; p=0.46, nested t-test; Figure 3A-C). However, the mIPSC amplitude was significantly higher in D2 mice compared to D2-controls (median(IQR); D2-control: 12.3(11.1-16.6) pA; D2: 18.9(14.4-23.8) pA; p=0.0092, nested t-test; Figure 3D&E). Thus, there appears to be an increase in quantal IPSC amplitude in the D2 dLGN, likely reflecting a post-synaptic up-scaling of inhibitory synaptic strength.

**Figure 2.**
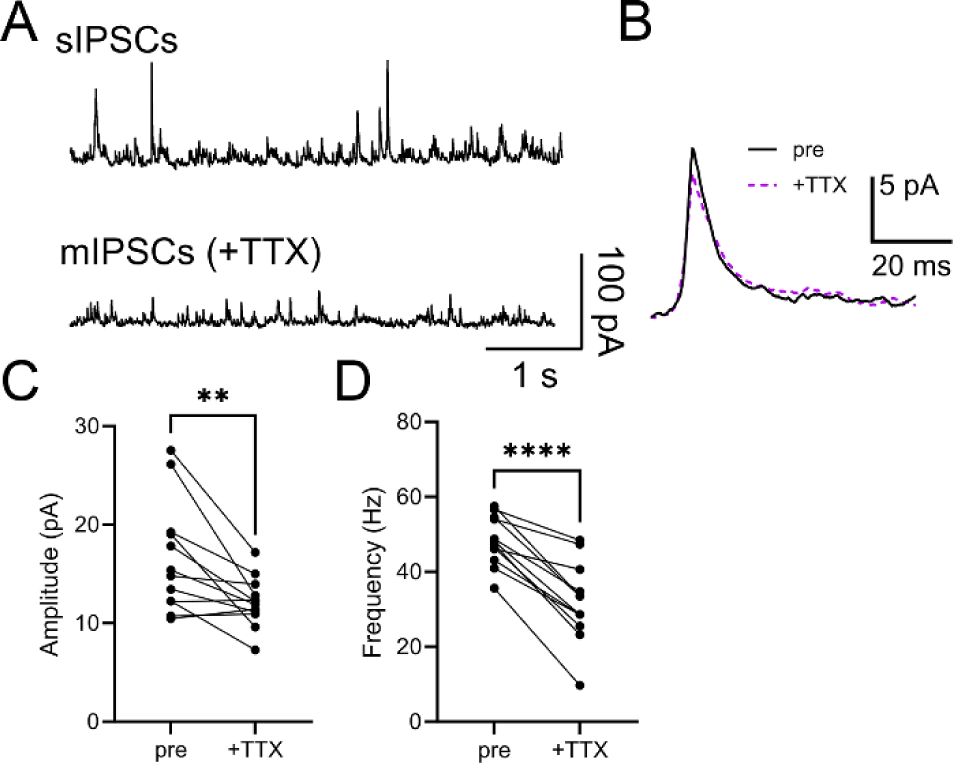
Acute blockade of voltage-gated Na+ channels alters inhibitory current properties. **A)** Example traces of inhibitory synaptic currents measured in the absence of stimulation before (sIPSCs) and after (mIPSCs) application of 0.5 µM TTX. **B)** Example mIPSC waveforms from the example in A. **C)** Group data showing that TTX application led to a significant reduction in mIPSC amplitude (**p<0.01, paired t-test). D) Group data showing that TTX application led to a significant reduction in mIPSC frequency (p<0.00001, paired t-test). n = 13 cells.

**Figure 3.**
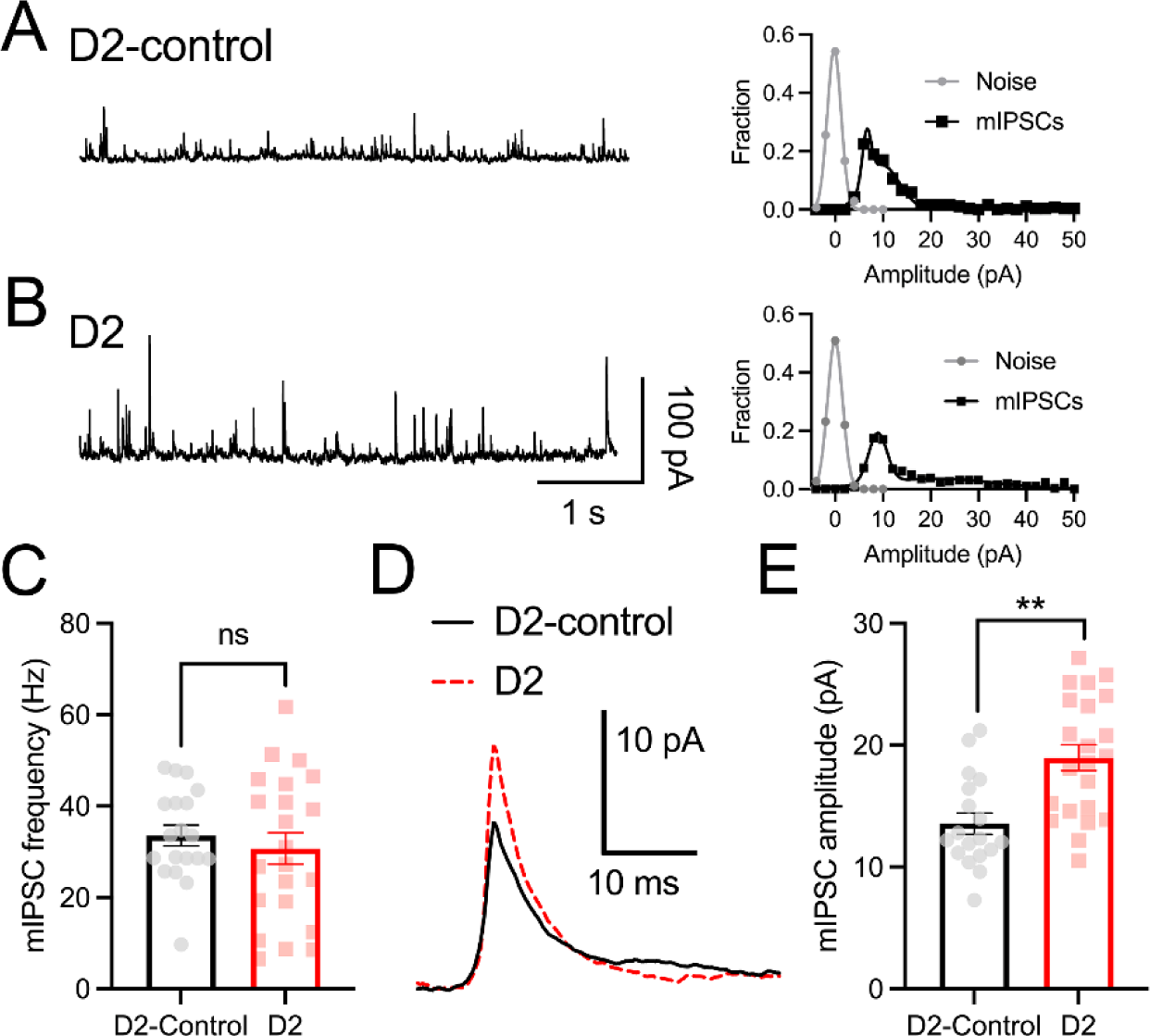
Elevation of quantal inhibitory synaptic current amplitude in dLGN brain slices from D2 mice. **A & B)** Miniature IPSC (mIPSC) recordings obtained while voltage-clamping at 0 mV in the presence of 0.5 µM TTX, 20 µM CNQX, and 50 µM D-AP5. Right, noise-amplitude histograms show good separation of measured mIPSC amplitudes from recording noise for the examples shown. **C)** Group data of mIPSC frequency of recordings from D2-control (19 cells from 8 mice) and D2 (22 cells from 9 mice). Nested t-test, ns p>0.05. **D)** Example average waveforms of mIPSCs detected from the recordings in A & B. **E)** Group data of mIPSC amplitude, **p<0.01, nested t-test.

We next applied two tests of synaptic scaling to our mIPSC amplitude data (Figure 4). In the first (Figure 4A-C), we used linear regression to fit the rank-order paired scatterplot of mIPSC amplitudes from D2-control and D2 recordings with a line (Turrigiano et al., 1998; Hanes et al., 2020). This produced a fit with a scaling factor (slope) of 1.45 and a y-intercept of –0.81 pA. We tested whether this linear regression was an accurate representation of the synaptic up-scaling by down-scaling the D2 mIPSC amplitudes by the 1.45 scaling factor, excluding those that fell below our mIPSC detection threshold (4.5 pA), and using a Komolgorov-Smirnov (K-S) test to determine whether the distribution of down-scaled D2 amplitudes was significantly different from the distribution of D2-control amplitudes. The cumulative distributions displayed some overlap but had a K-S test with p=0.0071, indicating statistically significant non-overlap of the two distributions (Figure 4B). When we examined the ratio of the rank-ordered D2 to D2-control mIPSC amplitudes as a function of D2-control amplitudes, we found considerable heterogeneity in the D2/D2-control amplitude ratios, rather than a consistent value (Figure 4C), which points toward non-uniform synaptic scaling.

**Figure 4.**
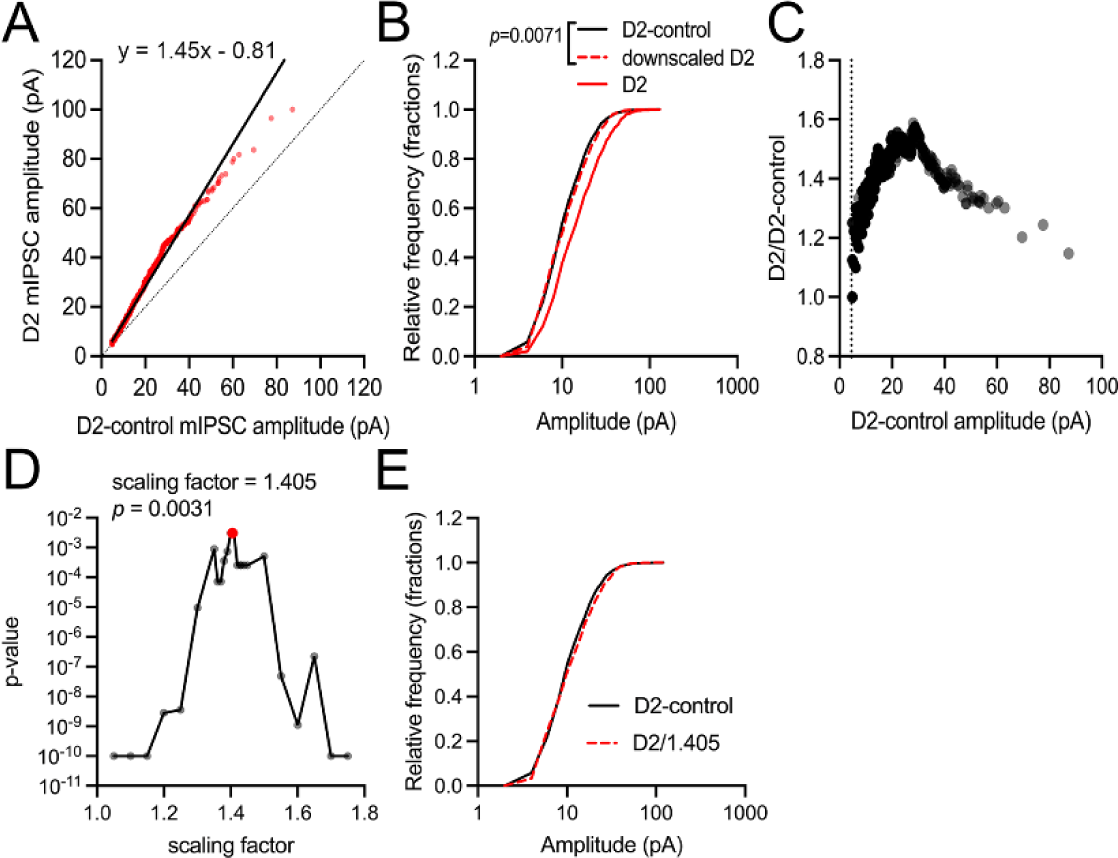
Tests for inhibitory synaptic scaling. **A)** Plot of rank-ordered mIPSC amplitudes from D2 and D2-control recordings (1900 mIPSCs from each group) fit using a linear regression model (solid line). Fit parameters had a slope of 1.45 and a Y-intercept of –0.81. Dotted line = unity. **B)** Cumulative histograms of mIPSC amplitudes from recordings of D2-control (solid black line), D2 (solid red line), and D2 that were downscaled using the 1.45 scaling factor from A. The downscaled mIPSC amplitude distribution significantly diverged from the D2-control distribution (p=0.0071, K-S test). **C)** Rank-ordered mIPSC ratio plot shows that D2/D2-control ratios do not converge at a single scaling factor. **D)** Iterative scaling approach. K-S test p-values from comparisons of the iteratively down-scaled D2 distribution plotted against scaling factors. The highest p-value was 1.405, with p=0.0031. **E)** Cumulative distributions of the D2-control mIPSC amplitudes and the D2 distribution down-scaled by 1.405.

For the second approach, we applied an iterative scaling strategy (Kim et al., 2012) by down-scaling the D2 mIPSC amplitudes by a range of scaling factors (1.05-1.75), excluding resulting values that fell below the 4.5 pA detection threshold, and comparing the resulting cumulative distributions with the D2-control mIPSC amplitude distribution using a K-S test (Figure 4D&E). In this case, a scaling factor of 1.405 produced a distribution with the highest K-S test p-value (p=0.0031), reflecting the closest alignment of the down-scaled D2 mIPSC distribution with the D2-control distribution. Together, these analysis results indicate that inhibitory input to dLGN TC neurons undergoes synaptic scaling by a factor of approximately 1.4-1.45 in D2 mice, although the range of D2/D2-control ratios and the observation that down-scaled D2 distributions significantly diverge from the control distribution point away from a uniform scaling of all inhibitory synapses on individual TC neurons. In the Discussion section, we elaborate on interpretation of this and the potential underlying mechanisms.

These above results point to an enhancement of inhibition at the post-synaptic GABA receptors on dLGN TC neurons. To test whether this represents a modulation of single channel conductance, as might occur by changes in receptor stoichiometry or modulation effects, versus a change in the number of synaptic GABA receptors, we performed a peak-scaled nonstationary fluctuation analysis of the mIPSC decay phase from recordings in D2-control and D2 TC neurons (Traynelis et al., 1993) (Figures 5 and 6). In the datasets used for this analysis, there was no significant correlation of mIPSC decay time constant with mIPSC amplitude, indicating adequate voltage clamp. We found that on average, D2-control inhibitory synapses had a single channel conductance (γ) of 18.8±2.4 pS (n=15 cells, 6 mice), which was not significantly different from measurements obtained from D2 TC neurons (17.2±2.5 pS; n = 15 cells, 6 mice; p=0.64, nested t-test). Synaptic GABA_A_ receptors on dLGN TC neurons are likely comprised largely of α_1_β_2_γ_2_ subunits based on immunostaining and *in situ* hybridization studies (Persohn et al., 1992; Wisden et al., 1992; Fritschy and Mohler, 1995; Pirker et al., 2000). Our measured single channel conductance values align closely with measurements taken from TC neurons in other thalamic nuclei which have a similar receptor complement (Browne et al., 2001; Schofield and Huguenard, 2007). Although we did not detect changes in γ, suggesting the composition of synaptic GABA receptors were unchanged in D2 mice, we did find that D2 TC neurons had a higher GABA receptor number (N) at each synapse (27±4) compared to D2-controls (15±1; p=0.02, nested t-test) (Figure 6C). Thus, an increase in the number of post-synaptic GABA receptors on dLGN TC neurons appears to contribute to synaptic scaling of dLGN inhibition in D2 mice.

**Figure 5.**
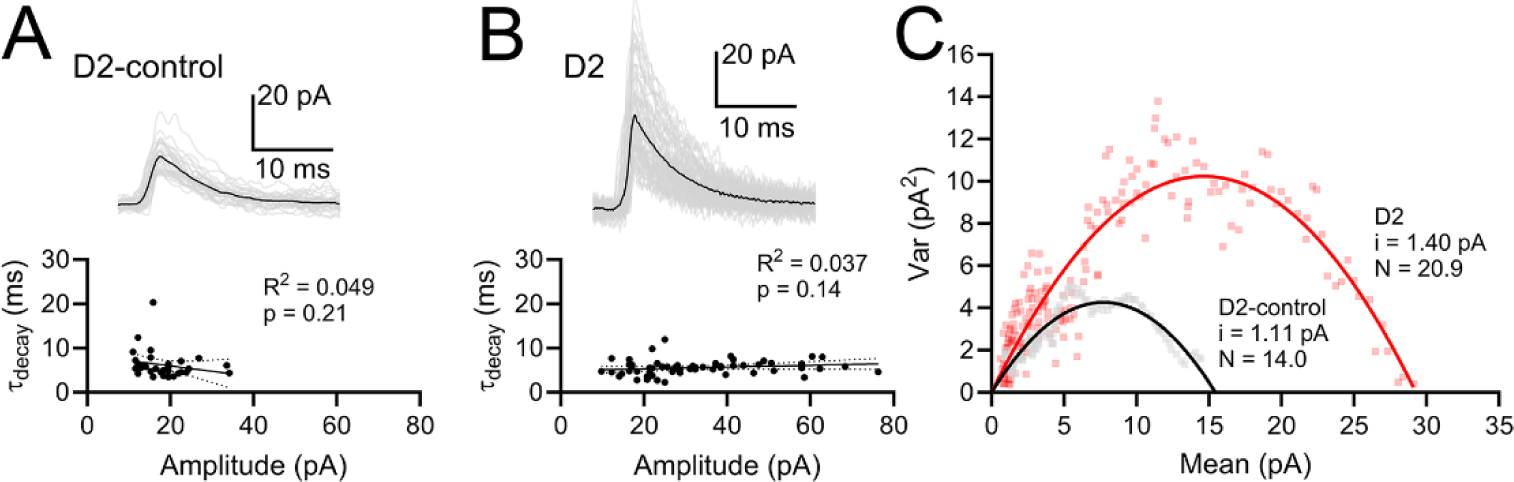
Peak-scaled nonstationary fluctuation analysis of mIPSCs. **A&B)** Example individual mIPSC events (gray traces) and mean event (black trace) from a D2-control recording (A) and D2 recording (B). Bottom, the mIPSC decay time constants were not significantly correlated with mIPSC amplitudes indicating adequate voltage-clamp. **C)** Variance-mean plot from examples in A and B, showing parabolic fit parameters including single channel current (*i*) and peak number of open receptors (*N*).

**Figure 6.**
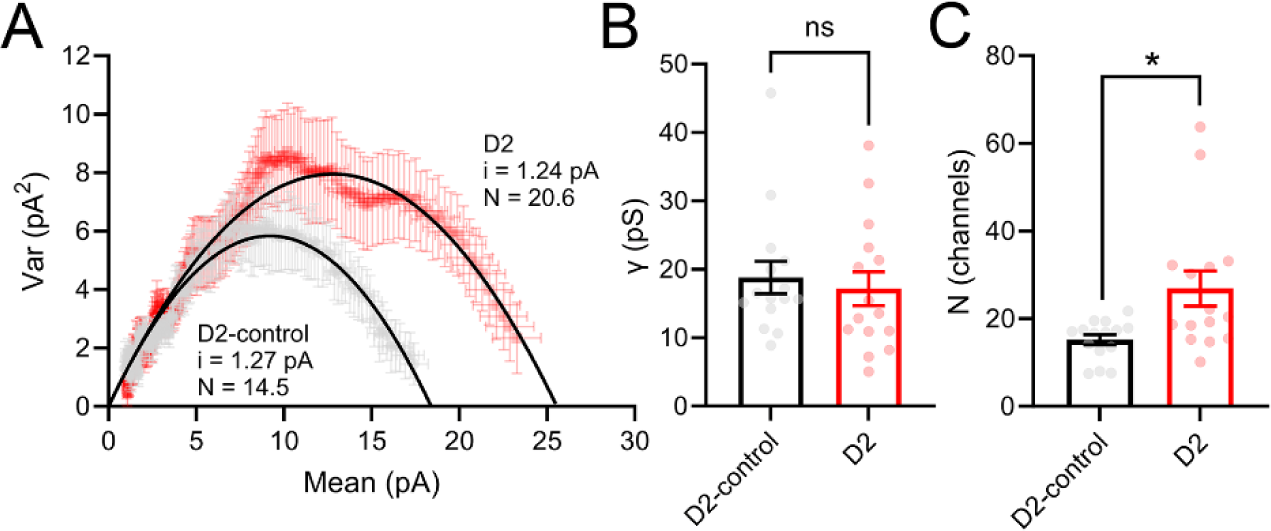
Peak-scaled nonstationary fluctuation analysis group data. **A)** Variance-mean plot of averaged (±SEM) variance and mean of mIPSC decay phases from D2-control recordings (15 cells from 6 mice) and D2 recordings (15 cells from 6 mice). Parameters of the parabolic fit are single channel current (*i*) and peak number of open receptors (*N*). **B)** Single channel conductance, measured by correcting for a 70 mV Cl^-^ driving force was not significantly different between D2-control and D2 (p>0.05, nested t-test. **C)** N was higher in D2 recordings compared to D2-control (*p<0.05, nested t-test).

To test whether we could detect increased GABA_A_ receptor expression in dLGN using an independent approach, we performed immunofluorescence staining for GABA_A_-α1 subunits in fixed dLGN sections from D2-control and D2 mice and analyzed the relative intensity of GABA_A_-α1 staining in optical sections representing the peak staining intensity from each slice (Figure 7). In this approach, we found that GABA_A_-α1 immunofluorescence intensity was higher in sections from D2 mice (33.7±1.5, n=17 mice) vs. D2-control mice (26.9±2.0, n=12 mice; p=0.0095, unpaired t-test). This result supports physiological data pointing to an increase in GABA_A_ receptor number at dLGN inhibitory synapses.

**Figure 7.**
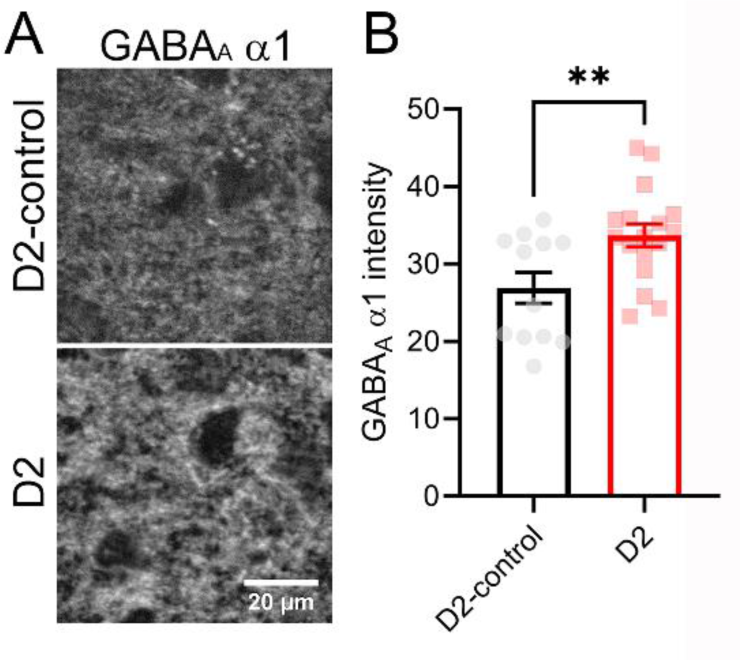
Increased GABA_A_-α1 immunofluorescence in dLGN of D2 mice. **A)** 2-photon confocal images of GABA_A_-α1 staining. Images are average intensity projections of 6 optical sections (0.3 µm spacing). **B)** Quantification of average pixel intensity shows an elevated GABA_A_-α1 staining in dLGN of D2 mice (p<0.01, t-test).

Finally, we sought to determine whether glaucoma led to disruption in the size and/or number of presynaptic GABAergic terminals in the dLGN, as changes to the presynaptic vGAT have been implicated in activity-dependent presynaptic homeostatic plasticity (De Gois et al., 2005; Prestigio et al., 2021) (Figure 8). To test this, we performed immunofluorescence staining of fixed dLGN slices for vGAT and quantified vGAT terminal size, density, and staining intensity. The size of detected vGAT puncta was similar in D2-control and D2 dLGN sections (D2-control: 1.06±0.03 μm^2^, n = 13; D2: 0.99±0.04 μm^2^; p=0.17, nested t-test). Additionally, the density of vGAT puncta was not significantly different between D2-control and D2 (D2-control: 66.7±3.2 puncta/1000 μm^2^; D2: 71.4±2.5 puncta/1000 μm^2^; p=0.25, unpaired t-test). The staining intensity of identified vGAT puncta was comparable between D2-control and D2 dLGN (p=0.51, nested t-test). Thus, there does not appear to be an effect of glaucoma on presynaptic GABAergic structures in the dLGN.

**Figure 8.**
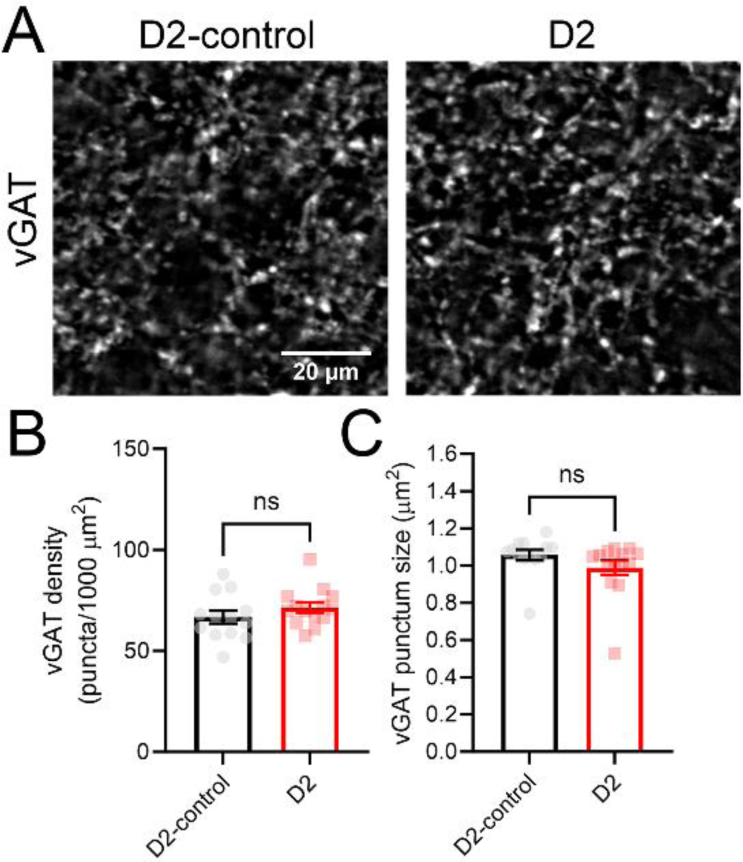
vGAT labeling is similar in D2-control and D2 dLGN. **A)** Example 2-photon confocal images (single optical sections) of vGAT labeling in dLGN. **B)** Quantification of detected vGAT punctum density (ns p>0.05, t-test; D2-control n = 13 mice; D2 n=15 mice). **C)** Quantification of detected vGAT punctum size (ns p>0.05, t-test).

## Discussion

The above results indicate that TC neurons in the dLGN of glaucomatous D2 mice show an increase in synaptic inhibition mediated, at least in part, through an upscaling of post-synaptic GABA_A_ receptors. We found that disynaptic feed-forward inhibition, evoked by optic tract stimulation, is diminished in D2 mice, although not as extensively as is expected if it simply reflected diminished drive from RG inputs. This points toward mechanisms that elevate inhibitory synaptic strength, perhaps compensating for diminished RG-to-LIN synaptic drive. Measuring quantal inhibitory currents (mIPSCs), we found that mIPSC amplitude was increased without detectable changes in frequency, indicating post-synaptic up-scaling of inhibitory synapses without alterations in inhibitory synapse number or vesicle release. A peak-scaled nonstationary fluctuation analysis indicated an increase in post-synaptic GABA_A_ receptor number without change in their single channel conductance and this was supported by staining showing enhanced staining for the major GABA_A_ receptor alpha subunit (GABA_A_-α1) in D2 dLGN tissue. Finally, analysis approaches testing for synaptic scaling point to an approximately 1.4-fold upscaling of inhibitory synapses, although those analyses revealed considerable non-uniformity in the extent of scaling across the inhibitory inputs to dLGN TC neurons, likely reflecting “divergent” scaling (Hanes et al., 2020; Koesters et al., 2024).

Homeostatic synaptic scaling is conventionally understood as a multiplicative process whereby all synapses of a type within a given neuron are scaled by uniform factor in response to altered activity (Turrigiano, 2012, 2017; Koesters et al., 2024). However, under many circumstances, scaling can be non-uniform, with synapses being scaled by divergent factors (Hanes et al., 2020; Koesters et al., 2024). Several analysis approaches have been used in attempts to distinguish these possibilities. These include testing whether the cumulative distributions of event amplitudes match after up- or down-scaling based on the linear fit parameters comparing rank-ordered control and experimental events (Turrigiano et al., 1998; Hanes et al., 2020, 2020). When we performed this analysis, we found that down-scaling our D2 mIPSCs by a scaling factor of 1.45 produced a distribution that had considerable overlap with the D2-control distribution. However, statistical comparison indicated quantitatively poor overlap suggesting that the D2 mIPSC amplitudes do not reflect a simple global multiplicative scaling process. Moreover, the peak K-S test p-value obtained using the iterative approach (Kim et al., 2012) only reached 0.0031 (at a scaling factor of 1.405), again indicating quantitatively poor overlap of the distributions. Finally, a plot of the ratio of rank-ordered D2/D2-control mIPSC amplitudes as a function of D2-control amplitudes showed a wide range of D2/D2-control ratios without clear convergence on a single value (Hanes et al., 2020; Koesters et al., 2024). Combined, these analyses point away from uniform multiplicative scaling of TC neuron GABA synapses. TC neurons receive the bulk of their inhibition from two sources: projections from the TRN and inputs from local interneurons, with LINs providing both dendrodendritic (F2 terminals) and axodendritic (F1 terminals) inputs to TC neurons (Bickford, 2019). mIPSCs recorded from TC neurons represent the combined input from each of these sources. The non-uniformity of scaling across the measured mIPSCs might reflect differential scaling at each of these synapse types or perhaps even differential scaling of synapses within a type. Whether this is the case, along with the underlying mechanism(s), remains to be determined.

At the ages used in this study (10-14 months), D2 mice have fairly minimal RGC somatic loss (Buckingham et al., 2008; Smith et al., 2022), although pre-degenerative processes such as RGC dendritic retraction and synaptic disassembly, diminished axoplasmic transport, and loss of RGC synaptic output in the brain are well underway (Crish et al., 2010; Calkins, 2012; Berry et al., 2015; Cooper et al., 2016; Smith et al., 2016, 2022; Risner et al., 2018; Van Hook et al., 2020). Within the dLGN, TC relay neurons also demonstrate clear structural and functional alterations including dendritic remodeling, somatic atrophy, altered intrinsic membrane properties, and enhanced action potential firing (Van Hook et al., 2020; Smith et al., 2022). Despite the diminished excitatory synaptic drive and altered intrinsic excitability, we have not seen evidence for excitatory synaptic scaling in D2 TC neurons. This might be tied to mouse age, as prior work has shown that silencing of retinal ganglion cell spiking or altered visual experience during a developmental critical period can alter excitatory glutamate receptors at retinogeniculate and corticothalamic excitatory synapses, respectively (Krahe and Guido, 2011; Louros et al., 2014).

While homeostatic synaptic scaling is thought to act on inhibitory and excitatory synapses in a coordinated fashion in response to altered activity levels, the triggers for scaling at each synapse type are different (Wenner, 2011; Gonzalez-Islas et al., 2023). This suggests that inhibitory synapses might be able to show signs of synaptic scaling without parallel changes in excitation, depending on the trigger. As shown in studies using primary cultures of hippocampal and cortical pyramidal neurons, excitatory scaling is a cell-autonomous process whereby altered somatic Ca^2+^ concentration serves as an error signal linking perturbations in neuronal spiking behavior to activity of the Ca^2+^/calmodulin-dependent protein kinase IV (CaMKIV) that drives homeostatic trafficking of postsynaptic glutamate receptors (Goold and Nicoll, 2010; Turrigiano, 2012; Joseph and Turrigiano, 2017). Inhibitory synaptic scaling, on the other hand, operates independently of CaMKIV (Ibata et al., 2008; Goold and Nicoll, 2010; Joseph and Turrigiano, 2017), indicating it depends on another mechanism, possibly even independently of cell-autonomous spiking (Wenner, 2011). Of note, other work has suggested that altered synaptic transmission itself triggers excitatory scaling independently of spiking activity (Fong et al., 2015), while cell-autonomous spiking activity scales inhibitory synapses (Gonzalez-Islas et al., 2023), both in contrast to the “conventional” understanding and suggesting a diversity of mechanisms that might depend on cell type and methods used to modulate activity.

Our data do not resolve what specific facet of glaucomatous pathology and associated changes in dLGN function trigger GABAergic scaling in the D2 dLGN. Nor do our data currently answer whether this results from network-wide effects or instead from cell-autonomous or synapse-specific processes. Activity can clearly modulate inhibitory synaptic strength; in work by Kilman and colleagues, activity suppression via TTX led to downscaling of synaptic GABA receptor number in primary cortical cell cultures, although it is unclear whether that was cell-autonomous, as network effects of TTX would also reduce spike-driven synaptic activity (Kilman et al., 2002). Brain-derived neurotrophic factor (BDNF) is a non-cell-autonomous trigger for homeostatic plasticity and has been shown to homeostatically influence inhibitory synaptic strength in hippocampal neuron primary cultures (Rutherford et al., 1998, 1998; Swanwick et al., 2006). In the retinogeniculate pathway, BDNF is produced by RGCs, transported along the optic nerve to their outputs and released, presumably in an activity-dependent fashion, where it can act on TrkB (tropomyosin receptor kinase B) receptors present on RGC axon terminals and TC neurons (Bennett et al., 1999; Avwenagha et al., 2006; Van Hook, 2022). BDNF signaling in the dLGN is likely disrupted in glaucoma due to diminished optic nerve transport and loss of RGC axon terminals (Pease et al., 2000), which could contribute to effects on inhibitory synaptic strength, although diminished BDNF signaling has generally been shown to diminish inhibitory synaptic strength (Swanwick et al., 2006; Wenner, 2011), in contrast to the enhancement we report here.

Another potential non-cell-autonomous mechanism involves tumor necrosis factor α (TNFα), a cytokine released by glial cells in response to injury and altered neuronal activity levels and can regulate post-synaptic AMPA and GABA receptors (Stellwagen and Malenka, 2006; Pribiag and Stellwagen, 2013; Heir and Stellwagen, 2020; Heir et al., 2024). The time course of activity deprivation appears to influence the effects of TNFα; prolonged activity blockade via TTX leads to TNFα release from glia, triggering excitatory synaptic scaling in culture (Stellwagen and Malenka, 2006) and acute TNFα application downregulates inhibitory synaptic function via both pre- and post-synaptic mechanisms (Pribiag and Stellwagen, 2013). Glaucoma has been shown to lead to reactive glial responses in dLGN (Tribble et al., 2021), and we have shown previously that optic nerve injury triggers changes in microglia morphology consistent with reactive microgliosis (Bhandari et al., 2022). Release of TNFα in the glaucomatous dLGN, either from activated microglia or astrocytes might be an important trigger of inhibitory synaptic scaling, but this possibility, along with whether it could have localized effects sufficient to trigger non-uniform scaling, remains to be tested.

Glaucoma is notorious for its relatively slow progression and asymptomatic presentation in early stages (Quigley and Broman, 2006; Tham et al., 2014; Weinreb et al., 2014). This has the potential of leading to late diagnosis, delaying treatment to a point where vision loss is irreversible. The GABAergic synaptic scaling we describe here might contribute to a vast array of adaptive processes that homeostatically preserve vision despite progressing pathology (Calkins, 2021). These include altered RGC excitability, changes in mitochondrial efficiency, glial responses, and changes in synaptic inputs and response properties of retinorecipient neuron populations in the brain (Calkins, 2021). Further study is needed to determine the cellular mechanisms of adaptation to glaucoma throughout the visual system and whether, when, and how the balance tips from sight-preserving adaptation to irreversible vision loss.

## Acknowledgements

This work was supported by a NIH/NEI grant R01EY030507 and a Career Advancement Award from Research to Prevent Blindness (RPB) and The Glaucoma Foundation to MVH. The funders had no role in study design, data collection, analysis, decision to publish, or manuscript preparation. The authors are grateful to Jennie Smith for performing intraocular pressure measurements.

## Author Contributions

MVH: Study design, data collection, data analysis, funding acquisition, manuscript preparation. SM: Data collection, data analysis, manuscript review and editing.

